# hipFG: High-throughput harmonization and integration pipeline for functional genomics data

**DOI:** 10.1101/2023.04.21.537695

**Authors:** Jeffrey Cifello, Pavel P. Kuksa, Naveensri Saravanan, Otto Valladares, Yuk Yee Leung, Li-San Wang

## Abstract

**Summary:** Preparing functional genomic (FG) data with diverse assay types and file formats for integration into analysis workflows that interpret genome-wide association and other studies is a significant and time-consuming challenge. Here we introduce hipFG, an automatically customized pipeline for efficient and scalable normalization of heterogenous FG data collections into standardized, indexed, rapidly searchable analysis-ready datasets while accounting for FG datatypes (e.g., chromatin interactions, genomic intervals, quantitative trait loci).

**Availability and Implementation:** hipFG is freely available at https://bitbucket.org/wanglab-upenn/hipFG. Docker container is available at https://hub.docker.com/r/wanglab/hipfg.

**Supplementary information:** Supplementary data are available as BioRxiv supplemental files.

## 1. Introduction

Genome wide association studies (GWAS) have successfully identified genome-wide significant variants and genes of interest with respect to the traits and disease under study (Uffelmann *et al*., 2021). However, understanding fully what these genetics signals mean biologically is still challenging. Post-GWAS analyses have benefited from recent advances in high-throughput sequencing technologies designed to characterize biology beyond the genome sequence, including epigenetic profiling, chromatin interactions, and gene-variant associations (Cano-Gamez and Trynka, 2020). Post-GWAS analyses, including variant fine-mapping or functional annotation, are often powered by genomic data querying and analysis-ready tools applied to these functional genomics (FG) data. However, within even a single assay or datatype of these FG data, there may be differences in file formats, included meta-information, and normalization standards. For example, in expression quantitative trait loci (eQTLs), the Genotype-Tissue Expression (GTEx) project (Aguet *et al*., 2017) published eQTLs in a 9-column format, and the eQTL catalogue (Kerimov *et al*., 2021) published the same reprocessed GTEx data with 19 columns. Integrating FG datasets into any analysis requires preprocessing steps specific to its source(s). Complicating this integration further, many data formats are not compatible with standard genomic tools used for querying such as bedtools (Quinlan and Hall, 2010), tabix (Li, 2011), and Giggle (Layer *et al*., 2018), therefore precluding them from immediate analysis following download. Some examples of common incompatibilities include:

- variable format and amount of provided meta-information across datasets and data sources,
- unconventional field names, unusual delimiters, or missing field name information,
- the use of different chromosome notations such as chromosome numbers or chrN chr-prefixed chromosome names,
- use of 1-based genomic coordinates instead of 0-based indexing in the UCSC Genome Browser BED style (Karolchik *et al*., 2003), or
- use of variable file formats such as text-based TSV formats not amenable to genomic querying.

Although workflows have been established to harmonize some individual data types such as GWAS summary statistics data (Lyon *et al*., 2021; Murphy, Schilder and Skene, 2021), no workflow yet exists to dynamically check, harmonize, and integrate diverse FG datasets and their metadata (Table S1).

Here we introduce the Harmonization and Integration Pipeline for Functional Genomics (hipFG), a robust and scalable pipeline for harmonizing FG datasets of diverse assay types and formats. hipFG can quickly integrate FG datasets for use with high-throughput analytical workflows, e.g., for analyzing current population-level studies such as UK Biobank (Sudlow *et al*., 2015) (500,000 individuals with >2,500 phenotypes). Fig S1 provides an example workflow for integrating hipFG with custom genomic analyses.

## 2. Material and Methods

hipFG includes datatype-specific pipelines to process diverse types of FG data. These FG datatypes are categorized into three groups: annotated genomic intervals, quantitative trait loci (QTLs), and chromatin interactions (Fig 1A). User-provided data descriptions (Fig 1B) guide hipFG in generating customized pipelines to harmonize input data and metadata. These pipelines include type-dependent and type-independent steps, checks, and corrections (Fig 1C, Fig S2).

**Fig 1:**
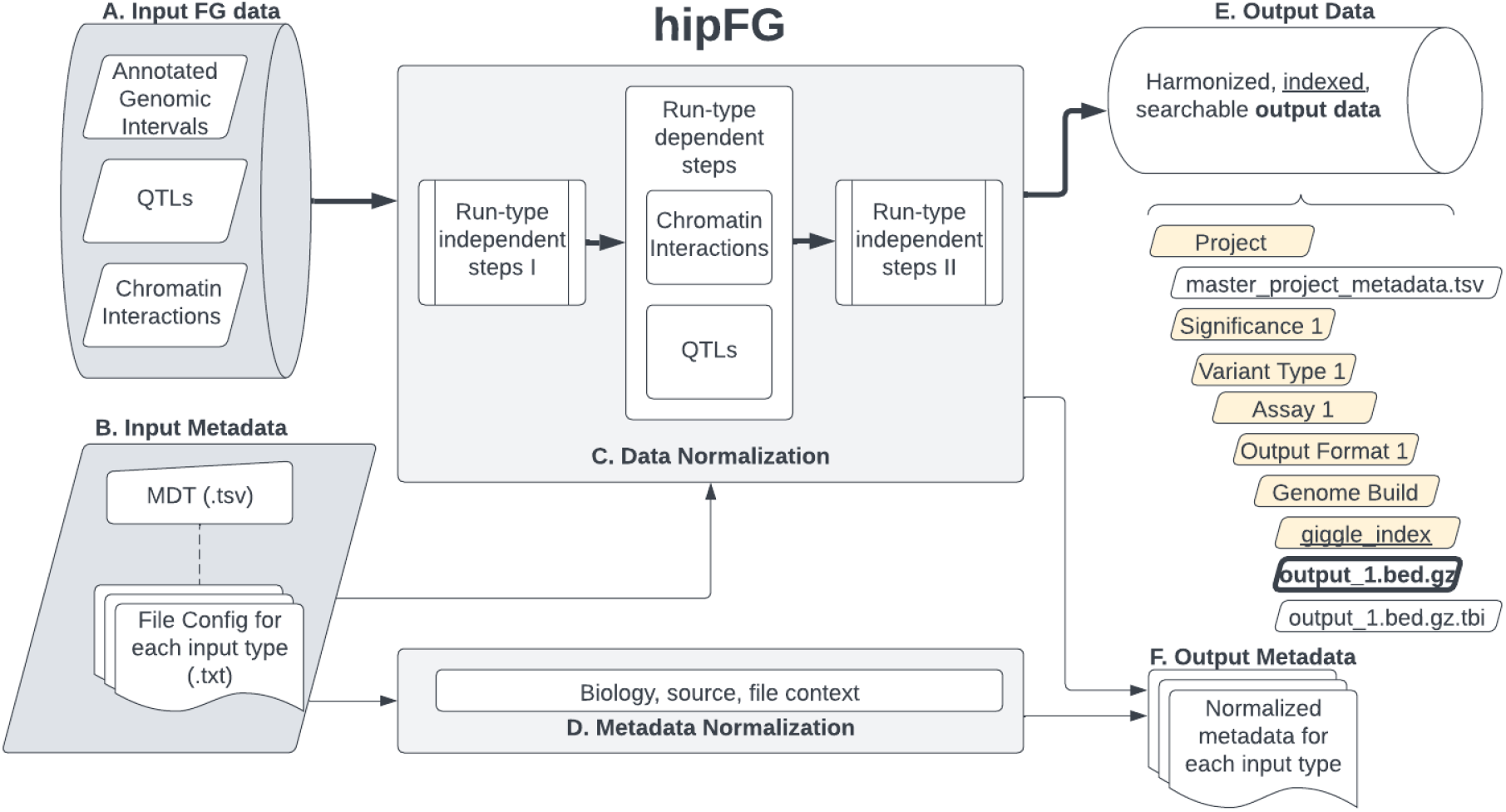
hipFG pipeline. Thin and thick lines indicate the flow of metadata and data, respectively. **A**. Input FG data can belong to any of these three groups. Non-standard file formats are acceptable. **B**. The input minimal descriptions table (MDT) provides biological, source, and file information describing all input FG data files. This includes paths to the file configs, which map provided columns to standard BED-format output fields. **C**. Data normalization includes steps dependent and independent of input data types (Details see Fig S2). **D**. Metadata normalization provides additional context to information provided in the MDT by providing additional biological context such as tissue and systems categories, recording provided publication information, calculating file attributes, and saving any supplemental information. **E**. The hipFG data outputs have standardized formats and are stored in an organized file-hierarchy according to metadata attributes. **F**. The output metadata contains the final output of the metadata normalization, integrated for all datatypes, and stored at the project level.

Type-independent steps are executed before and after the type-specific steps (Fig S2), and include genomic coordinate-based sorting, tabix-based indexing, metadata generation (Fig 1D), and consolidation of project-level metadata (Section 2.1). Type-dependent steps are highly variable and described in Section 2.2. Lastly, the output files are indexed for rapid query (Fig 1E) and organized alongside associated metadata (Fig 1F).

### 2.1 Metadata for input track data set

Data descriptions are provided via a minimal descriptions table (MDT) (Tables S2, S3) and an input-file configuration file (“file config”) (Fig 1B). The MDT includes biological (e.g., cell types, tissues), data source (e.g., DOI, project name), and file (e.g., data file path, paths to config files) information describing the inputs. The file configs (Fig 1B, Fig S3) associate input columns with standard output fields and describe other attributes of the input data, e.g., the existence of column names and 0-/1-based indexing. While the MDT (Table S3) describes all the input data files in a project, the file configs describe input-to-output reformatting for each data file type in the project.

To capture meta-information about the normalized output tracks, hipFG develops a structured, templated metadata table (Fig 1F, Tables S4,S5). After collecting initial metadata via the MDT, hipFG provides additional context to the provided input descriptions. Using biological annotations provided by the user according to open vocabularies derived from the FILER resource (Kuksa *et al*., 2022) hipFG assigns standardized “tissue” and “system” categories for each output file. Additionally, file information is calculated, including number of records/intervals, genomic base-pair coverage, and md5sum hash.

Any additional columns provided in the MDT are included in the “track description” metadata column in key-value pair format. This standardized metadata provides biological and source context to harmonized input data and any downstream genomic queries.

### 2.2 Standard FG data processing steps

The standard, type-independent steps are executed by the generated scripts regardless of output format and input datatype (Fig S2). One such standard step is column rearrangement according to the output BED-format, if necessary. Additionally, 1-indexed input data is adjusted to 0-indexing, and chromosome name formats are standardized. Lastly, the formatted output is sorted based on genomic coordinates, compressed via bgzip, indexed via tabix (Li, 2011), and indexed via Giggle (Layer *et al*., 2018) in its destination folder together with other tracks in the same category (Fig 1E). Each output file is matched with a hipFG standardized metadata (Fig 1F).

Standardized file outputs will be saved within a folder hierarchy in the following order, for applicable categories: association significance levels (by default, false discovery rate (FDR)<0.05 for QTLs), variant types (e.g., SNPs, insertions and deletions (INDELs)), assay types (e.g., ChIP-seq, RNA-seq, DNase-seq), output format types (e.g., bed19, bedInteract (Haeussler *et al*., 2018)), and genome builds (GRCh37/hg19 or GRCh38/hg38). This allows the outputs of projects with many datatypes and assays to be stored in an organized fashion as data collections, keeping similar datatypes together.

### 2.3 Type-dependent steps

hipFG’s processes inputs according to datatype-specific custom pipelines, with steps determined via file configs. The diverse input types allowed by hipFG necessitate separate consideration as described below.

#### 2.3.1 Annotated genomic intervals

Input files in BED format that include interval names, scores, and other fields but do not indicate a secondary target like QTLs or chromatin interactions do not require additional steps beyond the standard processing steps. The steps described in Section 2.2 are sufficient to standardize these types of data, which include narrow and broad peaks for epigenetic histone marks (ChIP-seq) or open chromatin regions (DNase-seq, ATAC-seq).

#### 2.3.2 QTLs

Expression, protein, or other types of QTL summary statistics data are highly variable in their format, names of provided fields, and which fields are included. QTL files therefore require additional processing steps (Fig S2C, QTL normalization). Following the type-independent steps (Section 2.2), allele correction and verification are carried out to normalize test statistics, allele frequencies, and variant IDs (to rsIDs when possible) (Fig S4). Additionally, QTL targets (e.g., genes, proteins) are annotated with GENCODE and UniProt when possible (Consortium, 2018; Frankish *et al*., 2018), providing gene symbol, strand, and distance from the transcription start site to the associated QTL. Lastly, if FDR is not provided, Benjamini-Hochberg *p*-value correction will be carried out per target gene-tissue pair. The final QTL results will be split by variant types (e.g., SNP and INDELs) and, if specified, significant QTLs (FDR<0.05 by default) will be extracted and saved separately from the full QTL summary statistics (Fig 1E).

#### 2.3.3 Chromatin Interactions

Chromatin interactions data (e.g., 3C, 4C, Hi-C (Dekker *et al*., 2002; van Berkum *et al*., 2010)) are processed such that both anchors (interacting sites) of each interaction are accessible by the genomic search tools mentioned above (Fig S2C, chromatin interactions normalization). With each interaction assigned a unique ID (numbered according to their original appearance), each interaction anchor is written on a separate line with its interaction ID plus an “A” or “B” to indicate it as the interaction source or target. The interaction “score” (integer in range [0-1000] indicating interaction strength) and “value” (a double calculated as the -log_10_ of the score), both expected in the UCSC “interact” format

(Haeussler *et al*., 2018), are calculated or assigned null values when missing. Additionally, each interaction is assigned a human-readable name combining the interaction’s data source, unique ID, score, and value. This name provides context to interactions when viewed with tools such as the UCSC Genome Browser (Karolchik *et al*., 2003). Lastly, any provided but unused columns are preserved as additional columns following the standard UCSC Interact Track Format fields (Haeussler *et al*., 2018).

## 3. hipFG usage

Following install, the hipFG.ini system configuration must be updated with the system-specific paths for the required programs, tools, and resources. The input MDT and file config(s) (Section 2.1, Fig 1B and Figs S2,3) are then sufficient for hipFG to process the input data (Fig 1A). hipFG can be executed with, at minimum, the MDT as its only argument. This generates (and executes) processing scripts for each of the input files and any number of input formats.

Lastly, the input files may be processed sequentially (on a single CPU or with multi-threading), or in parallel by chromosome-splitting or across input files. For full documentation, visit https://bitbucket.org/wanglab-upenn/hipFG.

## 4. Results

To demonstrate hipFG for FG data preparation/standardization and integration with high-throughput analysis workflows, we used hipFG to process and harmonize a heterogeneous, large-scale FG data collection including 109 eQTL catalogue eQTL datasets (Kerimov *et al*., 2021), 48 3DGenome chromatin interaction datasets (Wang *et al*., 2018), and 831 EpiMap samples with intervals annotated for one of 18 epigenetic states (Boix *et al*., 2021). These sources correspond to the three FG data types handled by hipFG, described in Sections 2.3.1-2.3.3, and include 17 billion variant-gene association records, 5 million genome-wide interactions, and 98 million annotated genomic intervals, respectively.

We then queried hipFG-processed and indexed data to annotate 10,823 genetic variants with genome-wide (*p*<5e-8) and suggestive (*p*<5e-6) significance from a recent Alzheimer’s disease GWAS (Bellenguez *et al*., 2022).

We found that variants were detected in a wide range of tissues within and across the three assay types (Fig S5). Following 1 MB binning based on tag variants for each chromosome, the genomic region chr17:45476979-46476979 contained 471 variant-tissue-assay combinations, the most of any such bin. This region includes known genes of interest including MAPT and KANSL1. These genes and this locus have been identified as sites of variants relevant to Alzheimer’s disease (Liu *et al*., 2015).

Additionally, hipFG’s processing speed was tested on five eQTL datasets from the eQTL catalogue dataset (Kerimov *et al*., 2021), demonstrating a linear run-time scalability with respect to input size (Fig S6). At the same time, all SNP reference alleles successfully matched to at least one genome reference, with an average 99.98% found in reference dbSNP b156 (Sherry *et al*., 2001), and the remainder resolved by matching against the GRCh38.p14 reference genome (Schneider *et al*., 2017).

In conclusion, hipFG enables scalable harmonization and integration of diverse, heterogeneous functional genomics data. Harmonized data and metadata allow for rapid querying and integration with high-throughput genetics workflows.

## Funding

This work was supported by the National Institute on Aging [U24-AG041689, U54-AG052427, U01-AG032984]; Biomarkers Across Neurodegenerative Diseases (BAND 3) (award number 18062), co-funded by Michael J Fox Foundation, Alzheimer’s Association, Alzheimer’s Research UK and the Weston Brain institute.

## Acknowledgements

We thank attendees of the American Society of Human Genetics Annual Meeting 2022 for valuable feedback on hipFG, as well as Luke Carter for testing hipFG installation.

## Notes

https://bitbucket.org/wanglab-upenn/hipFG/

### Competing Interest Statement

The authors have declared no competing interest.

## References

Aguet, F. et al. (2017) ‘Genetic effects on gene expression across human tissues’, Nature, 550(7675), pp. 204–213. Available at: https://doi.org/10.1038/nature24277.

Bellenguez, C. et al. (2022) ‘New insights into the genetic etiology of Alzheimer’s disease and related dementias’, Nature Genetics, 54(4), pp. 412–436.

van Berkum, N.L. et al. (2010) ‘Hi-C: A Method to Study the Three-dimensional Architecture of Genomes.’, JoVE, (39), p. e1869. Available at: https://doi.org/10.3791/1869.

Boix, C.A. et al. (2021) ‘Regulatory genomic circuitry of human disease loci by integrative epigenomics’, Nature, 590(7845), pp. 300–307. Available at: https://doi.org/10.1038/s41586-020-03145-z.

Cano-Gamez, E. and Trynka, G. (2020) ‘From GWAS to Function: Using Functional Genomics to Identify the Mechanisms Underlying Complex Diseases’, Frontiers in Genetics, 11, p. 424.

Consortium, T.U. (2018) ‘UniProt: a worldwide hub of protein knowledge’, Nucleic Acids Research, 47(D1), pp. D506–D515. Available at: https://doi.org/10.1093/nar/gky1049.

Dekker, J. et al. (2002) ‘Capturing Chromosome Conformation’, Science, 295(5558), pp. 1306–1311. Available at: https://doi.org/10.1126/science.1067799.

Frankish, A. et al. (2018) ‘GENCODE reference annotation for the human and mouse genomes’, Nucleic Acids Research, 47(D1), pp. D766–D773. Available at: https://doi.org/10.1093/nar/gky955.

Haeussler, M. et al. (2018) ‘The UCSC Genome Browser database: 2019 update’, Nucleic Acids Research, 47(D1), pp. D853–D858. Available at: https://doi.org/10.1093/nar/gky1095.

Karolchik, D. et al. (2003) ‘The UCSC Genome Browser Database’, Nucleic Acids Research, 31(1), pp. 51– 54. Available at: https://doi.org/10.1093/nar/gkg129.

Kerimov, N. et al. (2021) ‘A compendium of uniformly processed human gene expression and splicing quantitative trait loci’, Nature Genetics, 53(9), pp. 1290–1299. Available at: https://doi.org/10.1038/s41588-021-00924-w.

Kuksa, P.P. et al. (2022) ‘FILER: a framework for harmonizing and querying large-scale functional genomics knowledge’, NAR Genomics and Bioinformatics, 4(1), p. lqab123.

Layer, R.M. et al. (2018) ‘GIGGLE: a search engine for large-scale integrated genome analysis’, Nature Methods, 15(2), pp. 123–126.

Li, H. (2011) ‘Tabix: fast retrieval of sequence features from generic TAB-delimited files’, Bioinformatics, 27(5), pp. 718–719.

Liu, G. et al. (2015) ‘Identifying the Association Between Alzheimer’s Disease and Parkinson’s Disease Using Genome-Wide Association Studies and Protein-Protein Interaction Network’, Molecular Neurobiology, 52(3), pp. 1629–1636. Available at: https://doi.org/10.1007/s12035-014-8946-8.

Lyon, M.S. et al. (2021) ‘The variant call format provides efficient and robust storage of GWAS summary statistics’, Genome Biology, 22(1), p. 32. Available at: https://doi.org/10.1186/s13059-020-02248-0.

Murphy, A.E., Schilder, B.M. and Skene, N.G. (2021) ‘MungeSumstats: a Bioconductor package for the standardization and quality control of many GWAS summary statistics’, Bioinformatics, 37(23), pp. 4593–4596. Available at: https://doi.org/10.1093/bioinformatics/btab665.

Quinlan, A.R. and Hall, I.M. (2010) ‘BEDTools: a flexible suite of utilities for comparing genomic features’, Bioinformatics, 26(6), pp. 841–842.

Schneider, V.A. et al. (2017) ‘Evaluation of {GRCh38} and de novo haploid genome assemblies demonstrates the enduring quality of the reference assembly’, Genome Res., 27(5), pp. 849–864.

Sherry, S.T. et al. (2001) ‘dbSNP: the NCBI database of genetic variation’, Nucleic Acids Research, 29(1), pp. 308–311.

Sudlow, C. et al. (2015) ‘UK biobank: an open access resource for identifying the causes of a wide range of complex diseases of middle and old age’, PLoS medicine, 12(3), p. e1001779.

Uffelmann, E. et al. (2021) ‘Genome-wide association studies’, Nature Reviews Methods Primers, 1(1), p. 59. Available at: https://doi.org/10.1038/s43586-021-00056-9.

Wang, Y. et al. (2018) ‘The 3D Genome Browser: a web-based browser for visualizing 3D genome organization and long-range chromatin interactions’, Genome Biology, 19(1), p. 151. Available at: https://doi.org/10.1186/s13059-018-1519-9.

