## Supplementary Data for "hipFG: High-throughput harmonization and integration pipeline for functional genomics data"

### Supplementary Figures

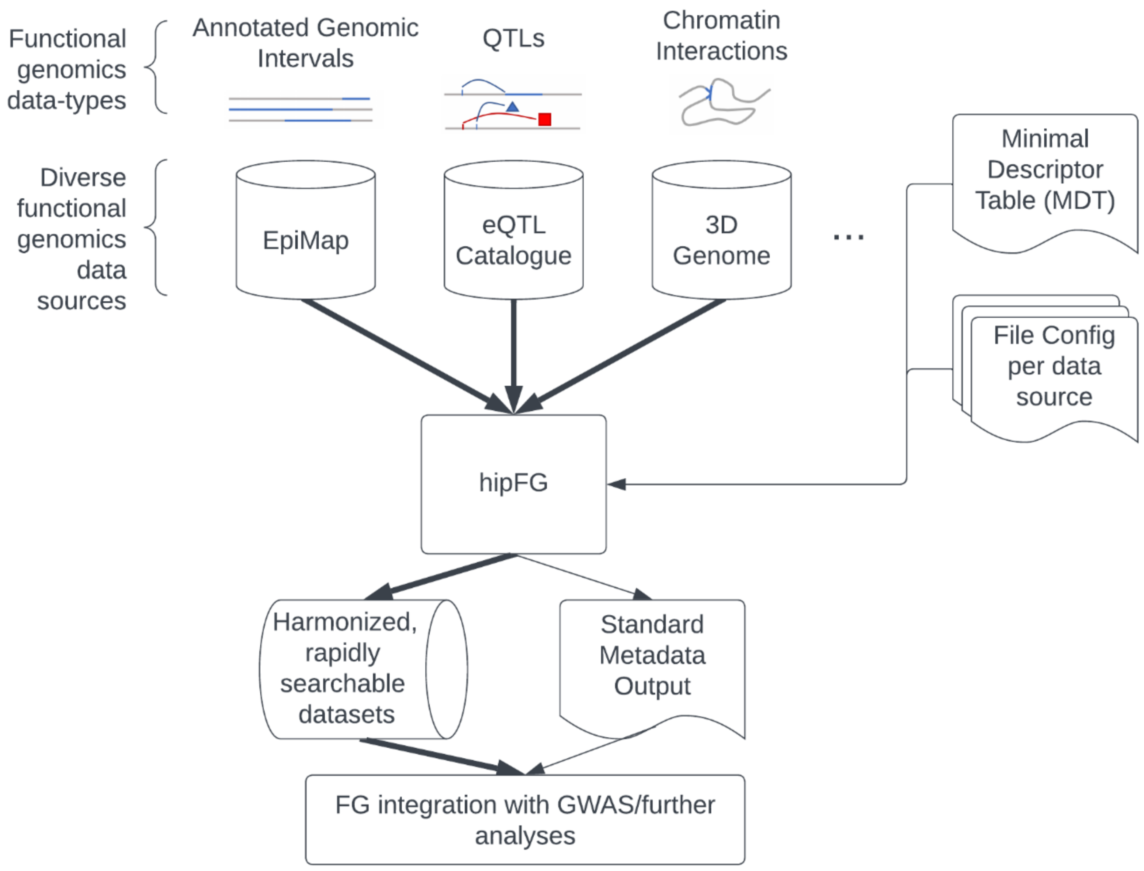


**Supplementary Figure S1.** High-throughput analysis workflow with hipFG. Input functional genomics data of diverse datatype and source are processed by hipFG into harmonized data and metadata. The resulting data and metadata are rapidly searchable and easily integrable into downstream workflows such as functional annotation of GWAS results, functional variant fine-mapping, colocalization or other analyses.

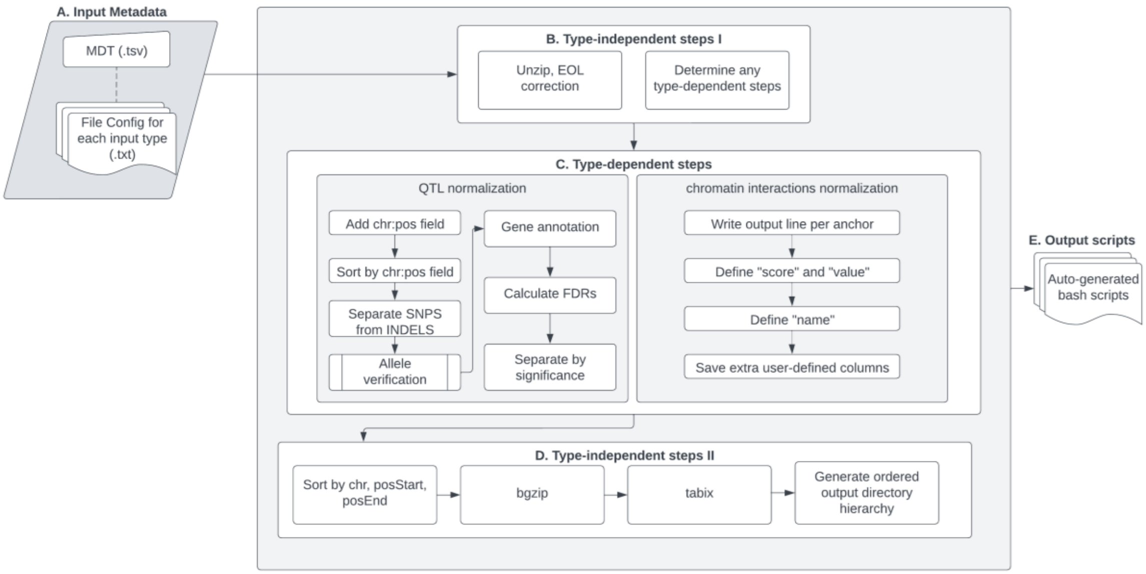


**Supplementary Figure S2:** hipFG data processing script-generation flow diagram. **A.** The input metadata is made up of a minimal descriptor table (MDT) *.tsv file containing biological, source, and file path information for all data files to be processed, as well as one or more file config (*.txt) files describing the input data format(s) and columns for each of data/file types. **B.** Steps to prepare/standardize input data regardless of datatype. **C.** hipFG normalization steps applicable to QTL and chromatin interactions data, written to pipeline bash scripts as needed. **D.** Steps to finalize output data regardless of datatype. **E.** The output pipeline bash scripts, with one script for each input file indicated in MDT.

**A.**

```
chr=chromosome
chrStart=position
chrEnd=
pval=pvalue
ref=ref
alt=alt
target=molecular_trait_id
beta_non_ref=beta
beta_se_non_ref=se
FDR=qval
tested_allele=alt
other_allele=ref
ac=ac
an=an
variant_id=rsid

CALC_AF=True
SPLIT_CHR=False
SEP_BY_SIGNIF=True
HAS_HEADER=True
IS_BASE_ONE=True
LOOKUP_COL=target
LOOKUP_TYPE=ensembl_gene_id
MULTIPLE_VARIANT_TYPES=True
```

**B.**

```
#DEFINE COLS
BED_FORMAT=bed9

##CONFIG INFORMATION
IS_BASE_ONE=False
ALL_INTERVALS_ONE=False
HAS_HEADER=False
```

**C.**

```
sourceChrom=sourceChr
sourceStart=sourceStart
sourceEnd=sourceEnd
targetChrom=targetChr
targetStart=targetStart
targetEnd=targetEnd
score=score_times_1000

IS_BASE_ONE=True
HAS_HEADER=True
```

**Supplementary Figure S3:** Examples of File Configs, as mentioned in Fig S2A, used to process **A**. eQTL catalogue (Kerimov *et al.*, 2021), **B**. EpiMap (Boix *et al.*, 2021), and **C**. 3D Genome (Wang *et al.*, 2018). The top section of each file config maps standard output fields on the left-hand-side to the provided fields in the input, on the right-hand-side. The use of BED_FORMAT in B. indicates to hipFG that all columns are provided with the correct names in the correct order (in a standard bed9 format in this case). Below the field assignment section is additional file information indicating to hipFG whether the genomic coordinates need to be corrected to 0-indexing, additional fields that need to be calculated by hipFG, and other context necessary for hipFG data normalization.

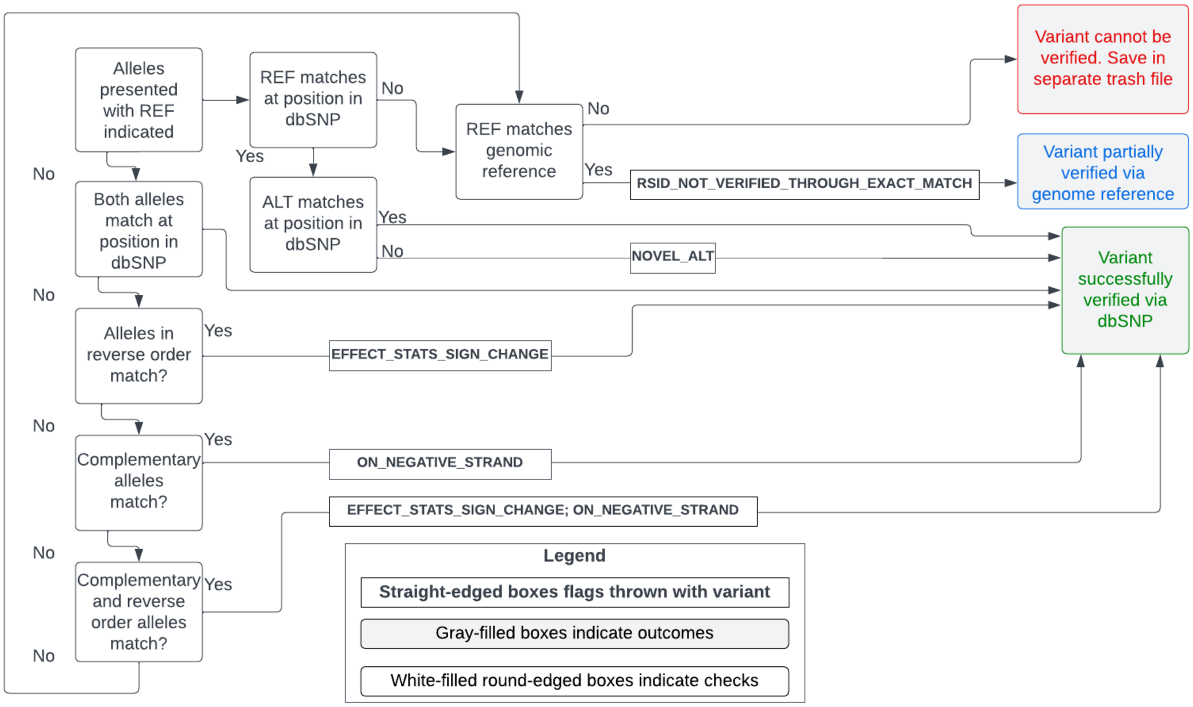


**Supplementary Figure S4:** hipFG allele correction flow diagram for processing QTLs. This flow diagram corresponds to the allele verification step of QTL normalization within Fig S2C. Provided alleles in the QTL data are matched to (by default) dbSNP (Sherry *et al.*, 2001) and, if this fails, the GRCh38.p14 reference genome (Schneider *et al.*, 2017). Any QC/normalization flags (shown as straight-edged boxes in the diagram) are added to the ‘QC_info’ key-term column of the final standardized QTL output.

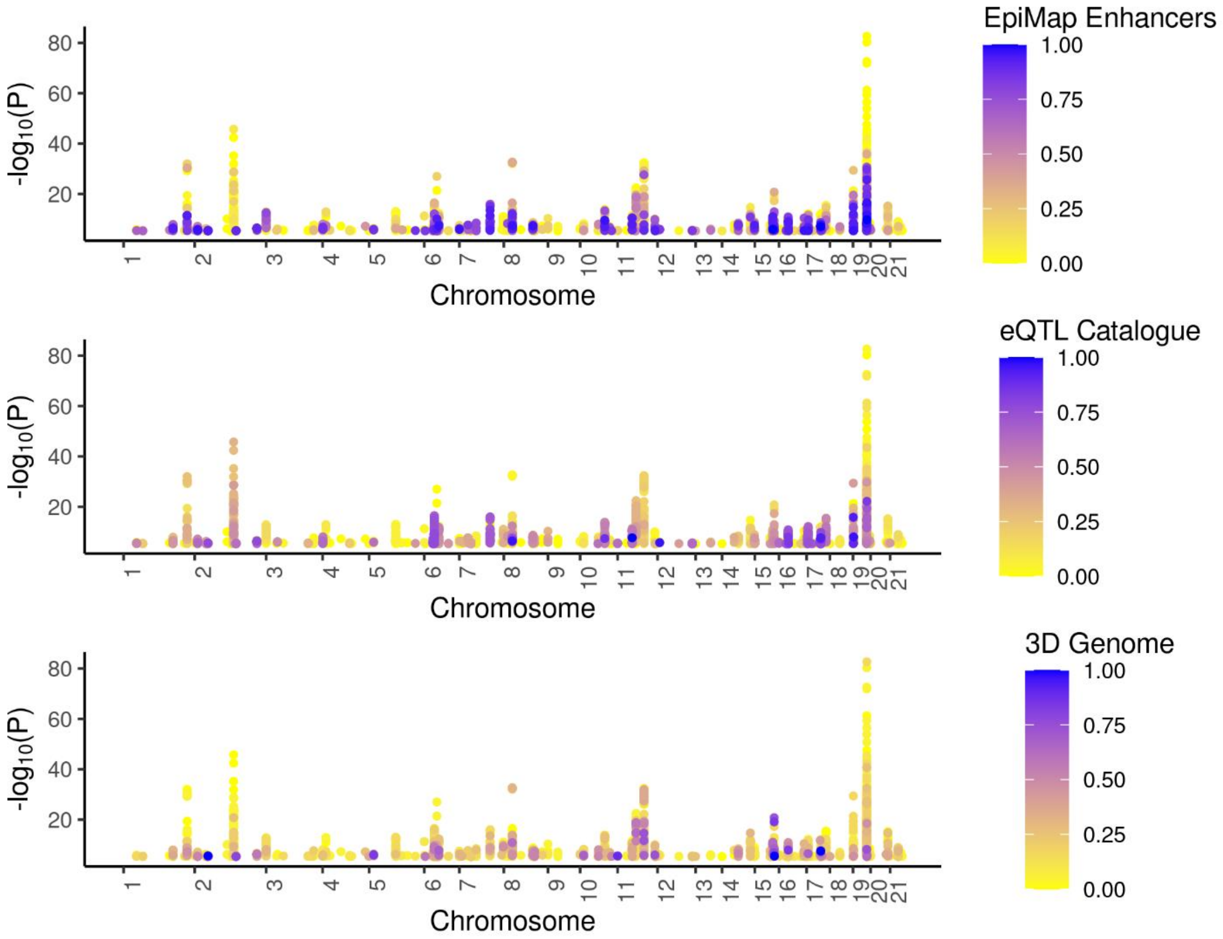


**Supplementary Figure S5:** We overlapped AD GWAS SNPs of suggestive significance (*p*-value < 5e-6) (Bellenguez *et al.*, 2022) with multiple FG datasets processed by hipFG. Fraction of tissue coverage by variants are shown in **A.** EpiMap Enhancers (Boix *et al.*, 2021), **B.** eQTL Catalogue (Kerimov *et al.*, 2021), and **C.** 3D Genome (Wang *et al.*, 2018).

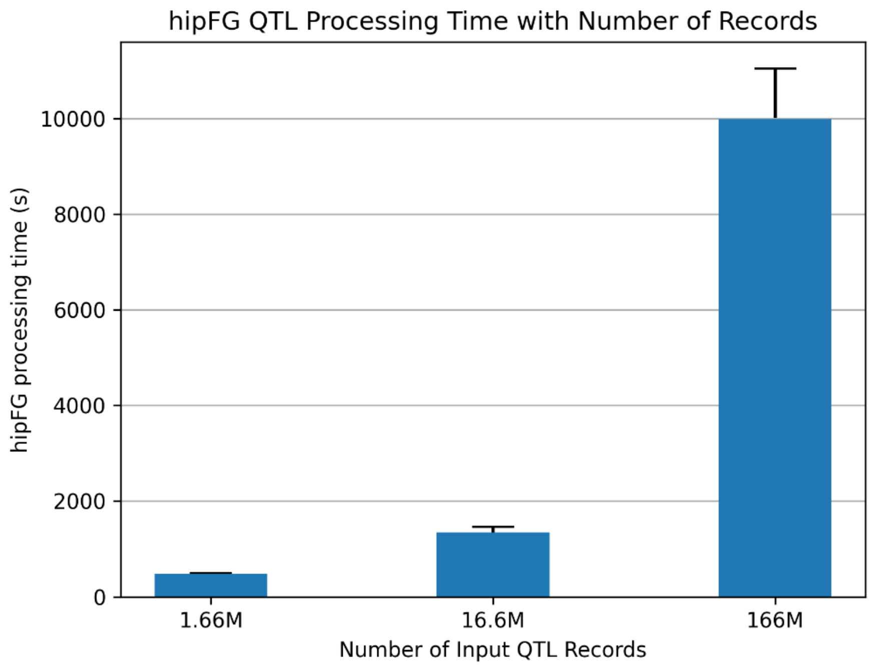


**Supplementary Figure S6.** hipFG running time scalability. hipFG processing time was measured on five example QTL datasets from eQTL catalogue (Kerimov *et al.*, 2021) that were subsampled at 1%, 10% or used in full (100%) and processed via hipFG. The average size without down sampling was 166 Million records, with an average processing time of 9,995 +/- 1,057 seconds. A 100x increase in input records resulted in a 21x increase in processing time, demonstrating linear scalability of hipFG. These examples yield an average QTL processing time of 60M QTL records per hour.

### Supplementary Tables

**Supplemental Table 1.** Summary of features for hipFG and other tools.

| **Features / Tool** | **FAVOR** (Zhou *et al.*, 2023) | **WGSA** (Liu *et al.*, 2016) | **MungeSumStats**(Murphy, Schilder and Skene, 2021) | **hipFG (this work)** |
|---|---|---|---|---|
| Scalable QC/standardization for multiple FG data types | No (variant annotations only) | | No (GWAS summary stats only) | ✓ |
| Cross-data source and cross-data type harmonization and integration of FG data and metadata | No (variants only) | | | ✓ |
| Standardized/unified (BED-based) indexed FG data formats | No (variants only) | | | ✓ |
| Standardized/unified metadata | | | | ✓ |
| Directly compatible/integration with standard downstream/annotation tools (bedtools (Quinlan and Hall, 2010), Giggle (Layer *et al.*, 2018)) | aGDS tools only | | ✓ | ✓ |
| Data categorization and organization/grouping | No (variant annotations only) | ✓ | | ✓ |
| Support for scalable FG data querying and organization/grouping | Yes (variants only) | | | ✓ |
| Support for scalable FG data querying and retrieval | Yes (variants only) | | | ✓ |

**Supplementary Table 2.** (Supplementary Tables.xlsx) Minimal descriptions table (MDT) fields. The MDT has 20 required input fields, which may be required for the hipFG scripts or for proper metadata characterization.

**Supplementary Table 3.** (Supplementary Tables.xlsx). An example MDT used to process 109 eQTL files from eQTL catalogue (Kerimov *et al.*, 2021). In this example, each row corresponds to one input file. However, all fields listed between output directory and project URL are shared across all input files.

**Supplementary Table 4.** (Supplementary Tables.xlsx) Output metadata table fields. The standardized output metadata table has 31 columns; some directly copied from the MDT and some generated or mapped throughout the hipFG run.

**Supplementary Table 5.** (Supplementary Tables.xlsx). An example standardized metadata output following running hipFG with the MDT provided in Table S4. This output has 31 output columns, compared to the 20 input columns of MDT. Splitting by variant type (SNV and other) and significance (significant, all tested) yields more output files than inputs.

**Supplementary References**


Bellenguez, C. *et al.* (2022) ‘New insights into the genetic etiology of Alzheimer’s disease and related dementias’, *Nature Genetics*, 54(4), pp. 412–436.

Boix, C.A. *et al.* (2021) ‘Regulatory genomic circuitry of human disease loci by integrative epigenomics’, *Nature*, 590(7845), pp. 300–307. Available at: https://doi.org/10.1038/s41586-020-03145-z.

Kerimov, N. *et al.* (2021) ‘A compendium of uniformly processed human gene expression and splicing quantitative trait loci’, *Nature Genetics*, 53(9), pp. 1290–1299. Available at: https://doi.org/10.1038/s41588-021-00924-w.

Layer, R.M. *et al.* (2018) ‘GIGGLE: a search engine for large-scale integrated genome analysis’, *Nature Methods*, 15(2), pp. 123–126.

Liu, X. *et al.* (2016) ‘WGSA: an annotation pipeline for human genome sequencing studies’, *Journal of Medical Genetics*, 53(2), pp. 111–112.

Murphy, A.E., Schilder, B.M. and Skene, N.G. (2021) ‘MungeSumstats: a Bioconductor package for the standardization and quality control of many GWAS summary statistics’, *Bioinformatics*, 37(23), pp. 4593–4596. Available at: https://doi.org/10.1093/bioinformatics/btab665.

Quinlan, A.R. and Hall, I.M. (2010) ‘BEDTools: a flexible suite of utilities for comparing genomic features’, *Bioinformatics*, 26(6), pp. 841–842.

Schneider, V.A. *et al.* (2017) ‘Evaluation of {GRCh38} and de novo haploid genome assemblies demonstrates the enduring quality of the reference assembly’, *Genome Res.*, 27(5), pp. 849–864.

Sherry, S.T. *et al.* (2001) ‘dbSNP: the NCBI database of genetic variation’, *Nucleic Acids Res*, 29(1), pp. 308–311.

Wang, Y. *et al.* (2018) ‘The 3D Genome Browser: a web-based browser for visualizing 3D genome organization and long-range chromatin interactions’, *Genome Biology*, 19(1), p. 151. Available at: https://doi.org/10.1186/s13059-018-1519-9.

Zhou, H. *et al.* (2023) ‘FAVOR: functional annotation of variants online resource and annotator for variation across the human genome’, *Nucleic Acids Research*, 51(D1), pp. D1300–D1311. Available at: https://doi.org/10.1093/nar/gkac966.
